# Disentangling gene expression burden identifies generalizable phenotypes induced by synthetic gene networks

**DOI:** 10.1101/2023.06.29.547078

**Authors:** Aqib Hasnain, Amin Espah Borujeni, Yongjin Park, Diveena Becker, Paul Maschhoff, Joshua Urrutia, Linus Rydell, Shara Balakrishnan, Yuval Dorfan, Christopher A. Voigt, Enoch Yeung

## Abstract

Large-scale genetic circuits are rapidly becoming critical components for the next generation of biotechnologies and living therapeutics. However, the relationship between synthetic and host gene expression is poorly understood. To reveal the impact of genetic circuits on their host, we measure the transcriptional response of wild-type and engineered *E. coli* MG1655 subject to seven genomically integrated circuits and two plasmid-based circuits across 4 growth time points and 4 circuit input states resulting in 1007 transcriptional profiles. We train a classifier to distinguish profiles from wild-type or engineered strains and use the classifier to identify synthetic construct burdened genes, i.e., genes whose dysregulation is dependent on the presence of a genetic circuit and not what is encoded on the circuit. We develop a deep learning architecture, capable of disentangling influence of combinations of perturbations, to model the impact that synthetic genes have on their host. We use the model to hypothesize a generalizable, synthetic cell state phenotype and validate the phenotype through antibiotic challenge experiments. The synthetic cell state results in increased resistance to *β*-lactam antibiotics in gram-negative bacteria. This work enhances our understanding of circuit impact by quantifying the disruption of host biological processes and can guide the design of robust genetic circuits with minimal burden or uncover novel biological circuits and phenotypes.

## Introduction

Genetic circuits are engineered biological systems composed of interacting genetic elements, such as protein coding sequences and promoters, that are designed to implement specified logics in living cells. The wide adoption of genetic circuits holds promise for significant advancements in biotechnology and medicine, offering new avenues for disease diagnosis and treatment, as well as providing sustainable methods for the production of biofuels, chemicals, and other valuable products. The use of these circuits can already be found in a range of tasks, including improving control of gene expression through directed evolution and high-throughput screens [1, 2, 3, 4], expanding biomanufacturing capabilities [5, 6, 7], advancing next-generation therapeutics and diagnostics [8, 9, 10], and sensing biotechnologically relevant compounds [11, 12, 13, 14].

Applications of synthetic biology in the environment and health require robust, predictable performance of gene circuits in the host organisms in which they reside. This is made challenging largely due to the synthetic construct imposing a burden on their host potentially resulting in reduced fitness, disruption of circuit performance, and undesirable and unforeseen phenotypes [15, 16, 17]. Though studies have been performed to measure the transcriptional response of the host under burden conditions [18, 19, 20], it is still unclear which synthetic construct components are driving the transcriptional disruption, especially in scenarios where burden is not realized through fitness defects (which have been shown to be highly correlated with cellular capacity [20]).

More than that, it is also not known how burden effects differ between constructs that are genomically integrated and constructs that are expressed from a plasmid. Quantitative methods to investigate genetic circuits have been developed to characterize and troubleshoot origins of failures [21, 22, 23], however methods that model the influence of the circuit on the host transcriptional response have been restricted to mathematical modeling and have not revealed how specific host gene expression is dysregulated [16, 24, 25].

Quantitatively disentangling the impact of synthetic gene expression on host gene expression is made challenging for several reasons. Firstly, synthetic genes are typically part of a network of genes, and if not as part of a network, at least an antibiotic resistance marker is required for screening transformants. Disentangling impact is also made difficult to confounders such as external inducers or the time point at which the measurements are taken. Lastly, distinct perturbations applied individually (e.g., two distinct synthetic genes) generally will not impact the host in an additive manner when applied simultaneously. Recent work has established machine learning for predicting the transcriptional response of combinatorial perturbations from single perturbations [26, 27]. Inspired by these works, we develop an adversarially-trained deep learning architecture [28] that allows for the prediction of unseen combinations of perturbations from transcriptomics data for data-driven modeling of gene expression burden.

In this paper, we seek to characterize and quantitatively model the transcriptional dysregulation of host biological processes that are perturbed by synthetic constructs. To characterize the impact, we analyze a set of *E. coli* MG1655 strains that have been engineered to express synthetic payloads integrated in the genome as well as from a plasmid. We examine significant changes in gene expression between engineered strains and the wild-type through differential expression analysis of RNA sequencing (RNA-seq) collected over 10 inducer combinations and 4 time points from each of 9 strains. This data are also used to identify 3 genes whose dysregulation explains differences in each strain’s stationary phase cell density. We train a classifier to identify 20 genes capable of distinguishing between wild-type and engineered strains and propose these as biomarkers of circuit engineering. Finally, our deep learning model is used to measure the influence of individual synthetic genes and combinations of synthetic genes on the host, identifying host genes which are disrupted, not by any single synthetic gene, but by a combination of synthetic genes. The burden imposed by synthetic constructs, which is not always reflected in a reduction in growth rate, was shown to be generalizable to new strains and new constructs through the phenotype of increased resistance to antibiotics in the *β*-lactam class, verifying the ability of our model to identify generalizable phenotypes induced by synthetic gene networks.

## Results

### RNA-seq characterization of a genomically-integrated genetic NAND circuit reveals significant disruption in transcriptome of E. coli

We sought to investigate the transcriptomic response of *E. coli* MG1655 engineered with genetic circuits in which the number of genetic components is modularly increased in size, ultimately resulting in a genetic circuit that performs Not AND (NAND) logic (Fig. 1A). The base circuit strain is comprised of four genes, driven by the constitutive pLacIq promoter [29], which switch regulatory states upon induction. In the final strain, *araC* and *lacI*, are used to implement control over the expression of two downstream repressors, *phlF* and *icaR*, which are activated in the presence of Ara and IPTG, respectively. The expression of yellow fluorescent protein (YFP) is controlled by the repressors to implement NAND logic, i.e. only in the presence of Ara and IPTG is YFP expression low, otherwise YFP expression is high (Fig. 1B). The resulting 9 strains used for this study are named wild-type (WT), landing pads, pBADmin, pTACmin, PhlF Gate, IcaR Gate, NAND Circuit, PhlF Gate (plasmid), and IcaR Gate (plasmid). Transcriptome profiles were generated for each strain, including the wild-type strain, across four time points: 5, 6.5, 8, and 18 hours post-induction, and 10 inducer conditions (Ara (mM)/IPTG (*μ*M): 0/0, 0/62, 0/75, 0/373, 6.2/0, 6.2/62, 7.5/0, 7.5/75, 37.3/0, 37.3/373) resulting in 783 wild-type and genomically-integrated profiles and 224 plasmid-based profiles for characterization. The time points were chosen to profile the transcriptome at early log phase, mid log phase, and stationary phase.

**Figure 1.**
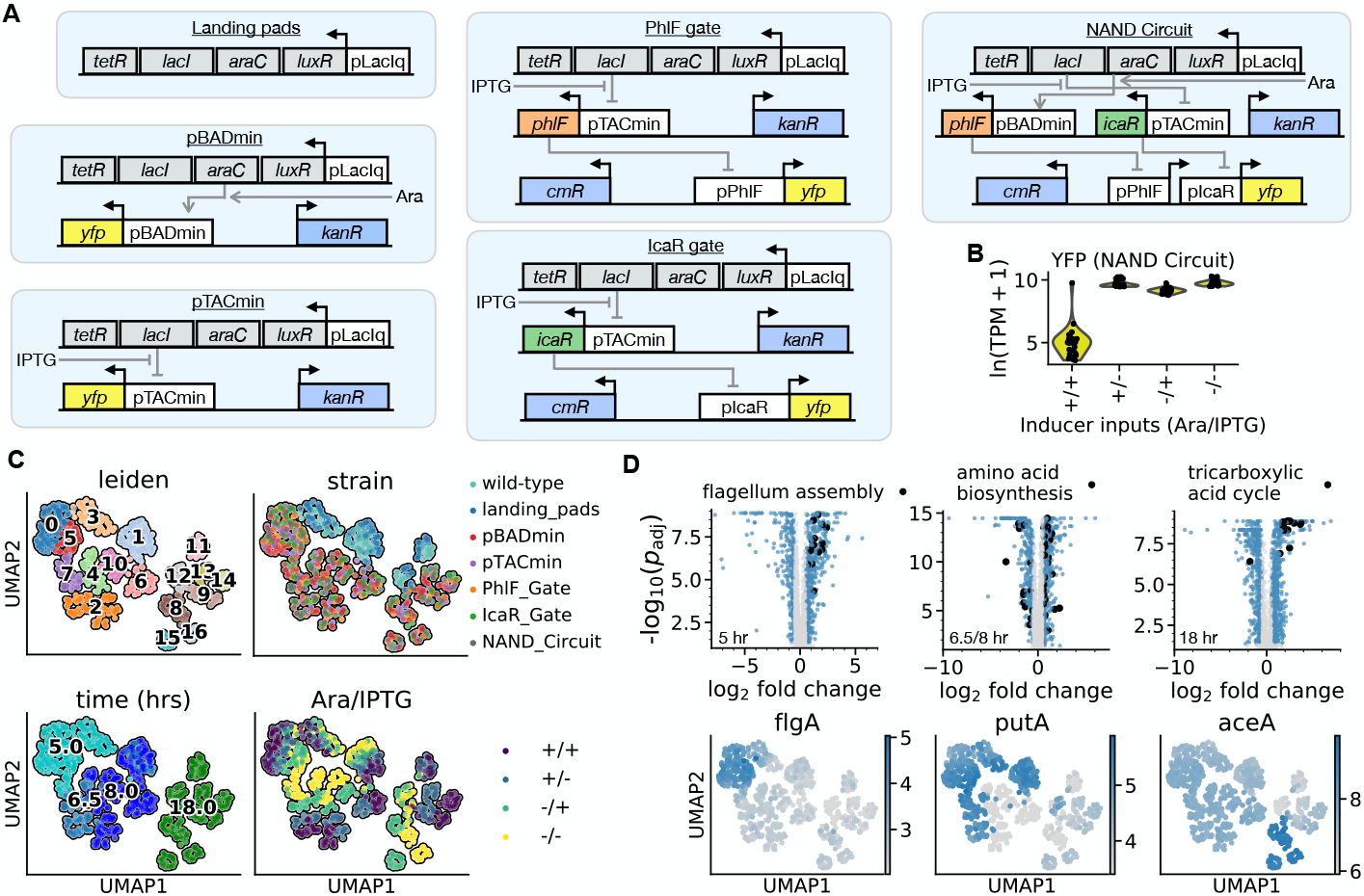
: Bottom-up construction and RNA-seq characterization of the genetic NAND circuit reveals significant disruption in *E. coli* genome. **(A)** Synthetic gene circuit schematics for the six strains used to characterize the impact of circuits on their host. **(B)** Distribution of the expression of YFP is shown for the genetic NAND circuit across the four inducer input states. Expression is quantified using *ln*(transcripts per million + 1). **(C)** Two-dimensional representation of wild-type and genomic integration circuit strain transcriptome profiles. The two dimensions correspond to the first two coordinates identified through graph-based clustering using PCA followed by UMAP. Graph-based clusters using the Leiden clustering algorithm are visualized in the top-left panel. The data points are colored by strain, time, and inducer combination in the top-right, bottom-left, and bottom-right panels, respectively. **(D)** Gene ontology enrichment analysis of differentially expressed genes identified flagellum assembly, amino acid biosynthesis, and tricarboxylic acid cycle to be over-enriched at 5, 6.5/8, and 18 hours, respectively. The upper panel shows, in large, black markers, the dysregulated genes belonging to the biological process in the legend. Genes marked in gray are not significantly differentially expressed. Genes marked in blue are differentially expressed, but do not belong to the biological process in the legend. The lower panels show marker genes from each biological process in the two-dimensional UMAP space. The color scale denotes the expression in ln(transcripts per million + 1).

To characterize the main sources of variation in the RNA-seq data, i.e. strain, timepoint, or inducer condition, we performed an integrated analysis of all 783 genomically-integrated and wild-type transcriptome profiles through graph-based clustering and manifold learning. Each profile consists of expression levels in *ln*(transcripts per million + 1), where transcripts per million (TPM) is calculated as described in [30], of 4,095 genes in the *E. coli* MG1655 genome. Distances between profiles can measure the level of similarity between two experimental conditions, however due to distances in high-dimensional space being untrustworthy [31], we first project the full transcriptome of each sample onto principal components of the data matrix. Next, to preserve both global and local structure of the manifold on which the data points lie, we perform Uniform Manifold Approximation and Projection (UMAP) to further reduce the gene-space to two dimensions for visualization [32, 33].

Graph-based clustering (Leiden clustering [34]) resolved distinct transcriptomic profiles, identifying 17 clusters which are characterized by combinations of strain, time, and inducer concentrations (Fig. 1C). We found that globally across clusters the main sources of variation are the strain and time point, while locally within a cluster the inducer combination dominates. The profiles are projected to two dimensions with UMAP for visualization purposes, and the Leiden clusters visually agree with variation in the reduced space.

The landing pads strain, containing four constitutively expressed heterologous genes and landing pads for integration elsewhere in the genome, is clustered together with the wild-type strain at all time points and inducer combinations. This strain is the closest to the wild-type *E. coli* as it does not contain any circuit components, except the four constitutively expressed genes responsible for sensing external inducers. Notably, the landing pads strain does not contain an antibiotic resistance gene. With the introduction of the antibiotic resistance gene, *kanR*, alongside YFP to construct the pBADmin and pTACmin strains, the similarity to the wild-type strain is diminished. This is consistent with previous proteomic studies that showed evidence of plasmid metabolic burden during fermentations is related to the production of antibiotic resistance gene products [35, 36].

Excluding the landing pads strain, the engineered strains’ transcriptome profiles are shown to be similar through clustering of pBADmin, pTACmin, PhlF Gate, IcaR Gate, and NAND Circuit strains at all timepoints and inducer combinations.

We next focused our attention to quantify the gene expression differences between engineered strains (excluding the landing pads strain) and the wild-type. We found 1846 genes to be statistically significantly differentially expressed when comparing the NAND strain to the wild-type strain; 767 after 5 hours, 683 after 6.5/8 hours, and 1138 after 18 hours. (Fig. 1D).

### Enrichment analysis reveals targeted disruption to genes associated with flagellum assembly, amino acid transport and biosynthesis, and TCA cycle

We found significant upregulation of flagellum assembly genes at the 5 hour time point when comparing engineered strains to the wild-type (Fig. 1D, left). Using Gene Set Enrichment Analysis (GSEA) [37, 38, 39], we identified that there is a 4.8-fold enrichment of flagellum assembly upregulation (Fisher’s exact test, FDR=4 × 10^−3^). *flgA*, required for the assembly of the flagellar P-ring formation, is upregulated 5.3 fold (Fig. 1D, left). Through GSEA we also found the enrichment of nitrate assimilation (4.6-fold enrichment, FDR=3 × 10^−3^), colanic acid biosynthesis (4.2-fold enrichment, FDR=1 × 10^−3^), aromatic amino acid biosynthesis (3.8-fold enrichment, FDR=4 × 10^−3^), anaerobic respiration (3.2-fold enrichment, FDR=2 × 10^−4^), and general transport associated biological processes (1.3-fold enrichment, FDR=2 × 10^−2^) 5 hours post-induction.

Significant down and upregulation of amino acid transport and biosynthesis was revealed through GSEA at the 6.5 and 8 hour time points (Fig. 1D, middle). Amino acid transport is enriched 3.3-fold (FDR=2 × 10^−2^) and amino acid biosynthesis is enriched 1.9-fold (FDR=4 × 10^−2^). The amino acid proline is catabolized by the proline dehydrogenase, *putA*, which is downregulated 11-fold (Fig. 1D, middle). Through GSEA we also found the enrichment of slime layer polysaccharide biosynthesis (5.8-fold enrichment, FDR=3 × 10^−2^), hexitol metabolism (5.3-fold enrichment, FDR=4 × 10^−2^), and cellular respiration (2.1-fold enrichment, FDR=3 × 10^−2^) 6.5 and 8 hours post-induction.

After 18 hours post-induction, GSEA reveals that the tricarboxylic acid (TCA) cycle dysregulation is enriched 7.7-fold (FDR=1 × 10^−6^) (Fig. 1D, right). This is consistent with the findings of proteomics and metabolic flux analysis which demonstrated an increase in TCA metabolic flux [36]. The upregulation ranges from 2.7-fold (*acnB*) to 13-fold (*aceA*) and one TCA cycle gene, *aspA*, is downregulated 3.5-fold. GSEA also reveals an 11-fold enrichment (FDR=4×10^−2^) in the glyoxylate cycle. This in conjuction with a 7.5-fold upregulation of *icd* suggests a shifting of carbon flux between TCA and glyoxylate cycles [40].

### Dysregulation of endogenous gene expression patterns explains fitness variation across genetically engineered strains

All engineered strains suffered from fitness defects observed through an increase in lag time and stationary phase density (Fig 2A). Strain pTACmin, different from pBADmin by a single promoter driving YFP expression, suffers from considerably greater defect in stationary phase density relative to other synthetic strains. This includes the NAND Circuit strain expressing three additional genes, one being a second antibiotic resistance gene which are known to be burdensome to the host [41]. This observed fitness defect in pTACmin is unexplained by the heterologous gene expression as strains with similar heterologous expression have different growth defects and strains with different heterologous expression have similar growth defects (Fig 2B). We then sought to explain this deviation in fitness in pTACmin through analyzing the response of host genes.

**Figure 2.**
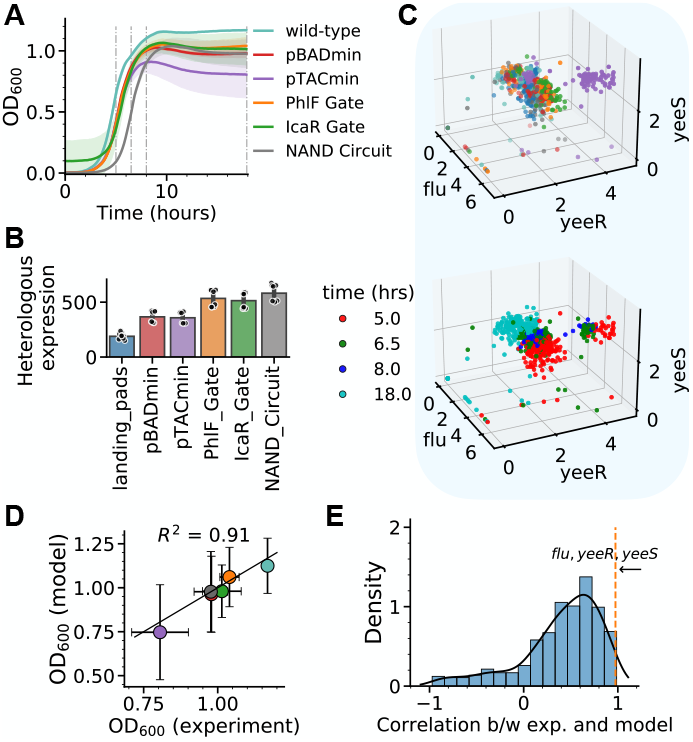
: Host gene dysregulation explains fitness variability across strains. **(A)** Growth curves for each strain used in the NAND circuit construction. Vertical gray lines denote the time points at which bulk RNA sequencing was performed on the individual strains. The averages and standard deviations were calculated using four biological replicates. **(B)** The total heterologous expression in each of the strains used in the NAND circuit construction where total expression is computed by summing the expression of all circuit genes within a replicate. The averages and standard deviations were calculated from eight replicates. **(C)** Expression (in ln(transcripts per million+1)) of three genes differentially expressed in pTACmin and not other circuit strains: *flu, yeeR, yeeS*, colored by strain (upper, see (A and B) for the color to strain key) and by time (lower). **(D)** The predicted stationary phase cell density (at 18 hours) of the six strains in (A), by the expression of genes *flu, yeeR, yeeS* shown in (C), are compared against their measured values (*R*^2^ = 0.91). The solid line depicts the *y* = *x* line. Experimental error bars represent the sample standard deviation in the measured cell density at 18 hours across four biological replicates. Model error bars represent the standard deviation over 192-fold cross validation. **(E)** Probability density of the correlation between mean predictions and mean stationary phase cell density obtained by randomly selecting 2000 triplets of genes. The correlation for the triplet of genes from (C) is indicated with a dashed orange line near *x* = 1.0.

We found a set of three genes to be consistently differentially expressed in the pTACmin strain when comparing to all other synthetic strains; the genes *flu, yeeR, yeeS* are upregulated 6, 3, and 2 fold, respec-tively, when compared to the pBADmin strain. The upregulation is present at 5, 6.5, and 8 hours regardless of inducer levels and diminishes at the 18 hour time point (Fig 2C).

*flu*, which codes for the protein known as Ag43, is responsible for colony form variation and autoaggregation in liquid media [42]. The activity of Ag43 can be switched between ON/OFF states through a competition between DNA methylation at GATC sites of the regulatory region of *flu* and repression by OxyR [43]. However, we find no evidence that levels of OxyR or DNA adenine methyltransferase (DAM) change between synthetic constructs (Supplementary Fig. 1). From a study investigating the role of Ag43 for biofilm formation, researchers showed that some Ag43 variants promote biofilm formation and result in a decrease in optical density (OD) [44]. *yeeR* and *yeeS* are co-transcribed as a single transcriptional unit along with *flu*, suggesting that they may as well play a role in biofilm formation [45]. In the pTACmin strain, correlations among *flu* and *yeeR* are significantly higher when compared to all other strains in this study, further suggesting a mechanism of co-regulation (Supplementary Fig. 2).

We next evaluated if the overexpression of *flu, yeeR*, and *yeeS* seen in the RNA-seq measurements can explain the fitness variability across strains. We constructed a linear model that predicts the optical densities at 18 hours, shown in Fig. 1C, from their transcriptome profiles at all time points (see Methods section “Predicting optical density from RNA-seq measurements”). The linear model is able to capture 91% of the variation in the optical densities across strains, using the expression of only these three genes (Fig 2D).

To test how statistically significant this result is, we compare to randomly selected sets of three genes and construct linear models for each random set, measuring their performance using the correlation between actual optical density and predicted optical density. From predictions of 2000 randomly selected triplets of genes, the probability density was empirically estimated through kernel density estimation (Fig 2E). From this distribution, we calculated the probability of achieving a higher correlation than that achieved by *flu, yeeR*, and *yeeS*. When comparing mean predictions and the mean measurements, this gene set achieved a correlation of 0.975. Of the 2000 tested gene sets, only two achieved a higher correlation with the maximum being 0.987. Empirically, the probability of randomly selecting a set of three genes that better explains the fitness defect is 2/2000 or 0.1%. This exceedingly unlikely probability of randomly selecting three genes to explain the fitness variability across strains, coupled with their dysregulation and role in biofilm formation, we conclude that the expressions of *flu, yeeR*, and *yeeS* are indicators of synthetic strain fitness.

### Time-course RNA-seq reveals a conserved set of synthetic cell state biomarkers indicative of circuit engineering

An integrated analysis of the RNA-seq data across wild-type and engineered strains show that the host transcriptome of engineered strains are transcriptionally similar to one another (Fig. 1C). This suggests the existence of host genes which are commonly dysregulated, giving rise to a synthetic cell state that, once characterized, can explain phenotypes resulting from construct burden and can be tuned based on the needs of an application. Moreover, knowledge of such synthetic-construct-induced genes can play a pivotal role in quantifying the burden imposed by expression of heterologous genes.

Ceroni et al. identified seven synthetic-construct-induced genes from RNA-seq measurements in *E. coli* transformed with plasmids that impose high burden on the host [19] as measured by a cellular capacity monitor circuit [20]. In their study, it was shown that the promoter from one of the synthetic-construct-induced genes, *htpG*, is burden-induced and a feedback controller was constructed to mitigate growth defects. We then wanted to see if the seven genes also are biomarkers of synthetic constructs when con-sidering strains with genomically integrated circuits. Due to the low copy number of genomically integrating synthetic constructs, the burden is thought to be lower than in the plasmid case.

We found a shared pattern of temporal dysregulation in the seven genes identified in [19] (Supplementary Fig. 3). When comparing the transcriptional profiles of the NAND strain to the wild-type strain, we find that a majority of these genes are not differentially expressed 5 hours post-induction. The genes then follow a pattern of upregulation at 6.5 and 8 hours followed by downregulation at 18 hours (and similar for the two plasmid strains). *htpG* is dysregulated no more than 2-fold, consistent with another study reporting plasmid-based host impact [21]. This then suggests that there are experimental contexts in which the seven identified genes are not necessarily burden-induced or indicative of engineered cell states. Motivated by this, we next sought to identify host genes that are indicative of circuit engineering, i.e. genes whose dysregulation is independent of the experimental conditions and depends only on possessing a synthetic construct or not. To identify these genes, i.e., genes whose dysregulation depends on the presence of a synthetic construct and not what is encoded on the construct, we trained a linear classifier (binary logistic regression; see Methods section “Classifying synthetic-construct-induced biomarker genes”) to predict, from a transcriptional profile, whether the sample belongs to a wild-type strain or to a engi-neered strain.

We found that most genes have insignificant contribution to classifying wild-type vs engineered strains from the transcriptional profiles. To quantify this, we compare the logistic regression (LR) coefficients and use them as a measure of feature importance (Fig. 3A). On a held-out test set of 504 out of 1007 total samples, the classifier correctly predicts the class of all 504 samples.

**Figure 3.**
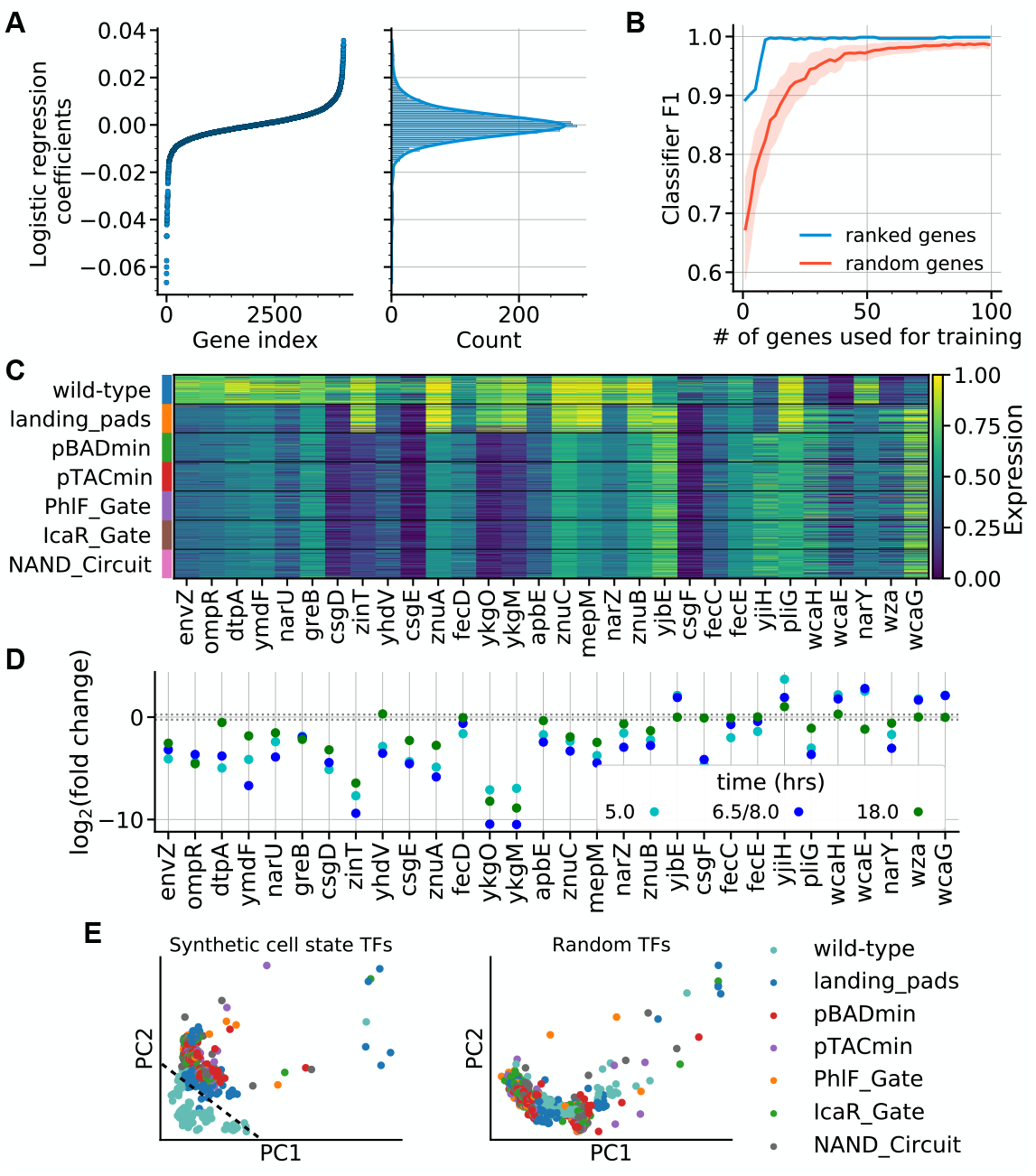
: The engineered cell state is classified by conditionally-independent, synthetic-construct-induced dysregulation. **(A)** Logistic regression was used to extract gene weights for classifying whether a transcriptional profile belongs to the wild-type or engineered strain class. The model coefficients are depicted on the left and a histogram of the coefficients on the right. **(B)** Logistic regression classifiers were trained on increasing number of genes, either ranked by the coefficients in (A) or by random selection and depicted here are the classifier F1 scores. **(C)** Expression (minmax normalized across all conditions) for synthetic cell state genes (30 genes with largest magnitude coefficient in logistic regression model) grouped by strain. **(D)** Fold change of the synthetic-construct-induced biomarkers in (C) comparing the NAND circuit to the wild-type condition. Horizontal dashed lines are displayed at ±0.25. **(E)** Two-dimensional representation of wild-type and genomic integration circuit strain whose transcriptome profiles are comprised of the 20 transcription factors associated with regulation of the synthetic cell state biomarkers (left) and 20 random transcription factors (right). The two dimensions correspond to the principal components which capture the largest amount of variance in the data. The black dashed line in the left panel indicates the linear separation between wild-type and synthetic strains.

Next, we quantified how well the coefficients of the LR classifier capture genes which are discriminative of wild-type and synthetic strains and found that on the order of 10 genes are needed for near perfect classification on the held-out test set (Fig. 3B). To do this, we re-trained LR classifiers on subsets of genes chosen by their magnitude of LR coefficients and used F1 score as the classification performance metric. Using this strategy, we find that using more than the top 10 genes for re-training provides diminishing returns for F1 score. Comparing this performance to randomly sampled genes for classification, we find that the identified important features significantly outperform random selection (Fig. 3B).

The 30 genes with largest magnitude regression coefficient are involved in disparate biological processes including nitrate assimilation, zinc ion transport, and sulfate transport (Supplementary Fig. 4). To showcase the conditionally-independent dysregulation across synthetic strains, we visualize these 30 genes in a heatmap shown in Fig. 3C. Here the RNA-seq data is grouped by strain, and within each strain are data from the four time points and 10 distinct inducer combinations. We see that regardless of the experimental context, these genes are dysregulated with striking fold changes ranging from 7 × 10^−4^ (*ykgM*) to 32 (*cysA*) (Supplementary Fig. 5). Of the 30 genes shown, 20 exhibit the similar transcriptional profiles in plasmid-based strains as in genomically integrated strains characterized in this study (Supplementary Fig. 6). We propose this set of genes as synthetic cell state biomarkers in engineered strains.

### Transcription factor regulation of engineered cell state biomarkers distinguish engineered from wild-type strains

To better understand interdependencies and coregulation between the proposed synthetic cell state biomarkers and to further validate the proposed set of biomarkers, we sought to identify shared transcriptional regulators, shared sigma factors, and gene essentiality. Two out of the 33 synthetic cell state genes are essential for growth in rich medium, *csgE* and *dtpA* [46], both of which are upregulated in synthetic constructs. *dtpA* plays a role in transport of di- and tripeptides [47] and is activated by the OmpR regulon [48] while *csgE* plays a role in curli production, assembly, and transport [49].

The housekeeping sigma factor, *σ*^70^, regulates the transcription initiation of the majority of engineered cell state genes (Supplementary Table 1). We also found that 5 out of 30 of the biomarkers are regulated by the stationary phase sigma factor, *σ*^38^. At the 18 hour time point the expression of *σ*^70^ is higher in the wild-type strain and the expression of *σ*^38^ is lower in the wild-type strain compared to engineered strains, suggesting an increased or early-induced stress response in engineered strains (Supplementary Fig. 7).

We found that the transcription of the engineered cell state biomarkers are regulated by 20 unique transcription factors (TFs) (Supplementary Table 2). To understand how significant the expression of the identified 20 TFs are for distinguishing engineered cell states from wild-type cell states, we measured how distinct they are through linear dimensionality reduction on only these 20 TFs. Using principal component analysis, we computed the directions of maximal variance in the data (subsetted to the 20 TFs) and projected the data onto these directions. We similarly followed this procedure when 20 randomly selected TFs are chosen instead of the identified 20. We found that when considering the 20 identified TFs the distance between wild-type and synthetic strains in the reduced space is larger than when considering 20 random TFs (Fig. 3E). Engineered strains are nearly linearly separable from the wild-type when considering only the 20 identified TFs, whereas there is no clear separability for the random TF case.

### A deep learning model quantifies the targeted impact of genetic circuits on the host transcriptome

The transcriptional processes of engineered microbes are heavily disrupted by the introduction of synthetic payloads. Though studies have shown that kanamycin resistance genes have significant effects on gene transcript levels in *E. coli* [41], there is no quantitative model that demonstrates this effect. To quantify the effects of individual synthetic construct elements on the host transcriptome, we develop a deep learning model, influenced heavily by prior architectures for deconfounding batch effects and other co-variates in transcriptomics data [50, 27], which disentangles the influence of individual perturbations on the host transcriptome.

Our deep learning architecture consists of a set of three autoencoders, one for gene compression and reconstruction, one for continuous covariate (perturbation) compression and reconstruction, and one for discrete covariate compression and reconstruction. Alongside are two discriminators, a perturbation discriminator and a covariate discriminator (Fig. 4A). Gene expression vectors representing transcriptomic profiles, *x*, are compressed by a variational autoen-coder to a latent state *z* which is distributed according to a standard normal distribution. The associated perturbations, *p*, and covariates, *c*, are compressed deterministically to latent states *q* and *d* of the same dimension as *z*. To facilitate the gene latent state, *z*, to be free from information about the corresponding perturbation and covariates, during training the encoders are penalized for producing latent states from which discriminators are able to predict the perturbations and covariates. In this way, *z* becomes a perturbation-free (and covariate-free) latent state. The influence of perturbations and covariates on the host genes are then captured in the latent space through a linear model of the form

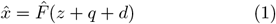

where 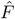 is the decoder tasked with predicting 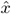, the reconstruction of the gene expression *x*. See the Methods section”Circuit-impact model” for details on model training, hyperparameter selection, and model evaluation.

**Figure 4.**
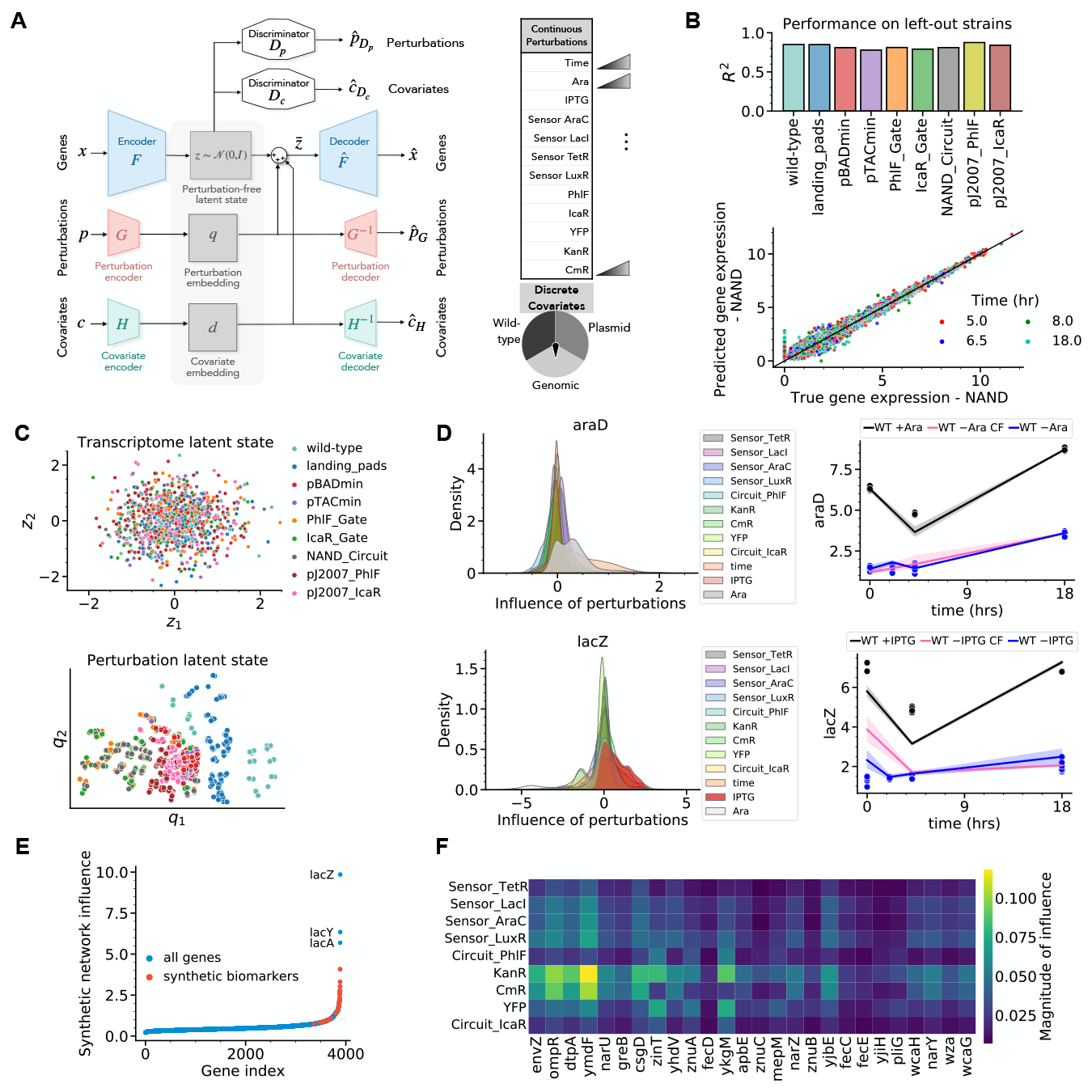
: Deep learning captures host gene dysregulation in unseen circuit combinations. **(A)** A schematic depicting the deep learning architecture for disentangling the influence of individual synthetic construct elements. The network consists of three autoencoders (and two discriminators) which transform inputs *x, p*, and *c* into latent states *z, q*, and *d*, respectively, and where *x* denotes a vector of genes, *p* denotes a vector of continuous perturbations and *c* denotes a vector of discrete covariates The list of perturbations and covariates are provided in the tables on the right. The discriminators are each tasked with predicting either the perturbation vector or the covariate vector. The gene encoder is penalized for producing latent states *z* from which the perturbations and covariates are able to be discriminated. **(B)** (Top) Results of leaving one strain out from training and testing on the left-out strain are measured through the coefficient of determination, *R*^2^. The results shown here are from the model selected via hyperparameter optimization. (Bottom) Predicted vs. true gene expression of the NAND Circuit strain from the circuit impact model trained on all other strains. **(C)** The first two-dimensions of the transcriptome latent state, *z*, is depicted for all samples *x* in the RNA-seq dataset. Below is the resulting two-dimensional perturbation latent state, *q*. All data points are colored by strain. **(D)** (Left column) Kernel density estimates of the influence of each continuous perturbation in *p* on the host transcriptome *x*, i.e. *∂x*_*i*_*/∂p*_*j*_, for two genes *araD* and *lacZ*. (Right column) Counterfactual experiments capturing the ability of the model to 1) simulate time responses to doses of inducers, and 2) recapitulate activity of known sugar utilization pathways. **(E)** Total influence of the synthetic genetic network on host genes. Influences on synthetic cell state biomarkers are visualized separately. **(F)** Individual influence of genes in the synthetic gene network on the synthetic cell state biomarkers.

We found that our circuit impact model accurately captures the impact of perturbations on unseen strains, achieving an *R*^2^ of over 0.75 on all left-out strains (Fig. 4B, top panel). This includes the NAND Circuit strain in which all synthetic regulatory interactions are present, demonstrating that the deep learning approach captures the compositional impact of regulatory elements. Specifically, the model captures host gene dysregulation upon composition of both *icaR* and *phlF* in the NAND Circuit strain, after only being trained on the impact of each individually (Fig. 4B, lower panel). Here we have also included two plasmid-based strains, pJ2007 PhlF, and pJ2007 IcaR for which RNA-seq measurements were also taken across IPTG/Ara doses and time points.

Once the model is trained, the latent state *z* no longer carries information of the perturbations and covariates (Fig. 4C, top panel). This is enforced through adversarial training and also through enforcing that *z* follow a standard normal distribution. Thus, the effects of perturbations and covariates can be fully captured in the latent space. Our model also provides an interpretable perturbation latent state that can be used to quantify the similarity and differences between the impact of perturbations across strains (Fig. 4C, bottom panel.) The first dimension of the perturbation latent state, *q*_1_, captures the variation of perturbations across strains while the second dimension, *q*_2_, captures the temporal perturbation differences (Supplementary Fig. 8).

Testing to see if our model has captured known biological mechanisms of regulation, we found a positive influence of the *araBAD* operon by arabinose and a positive influence of the *lacZYA* operon by IPTG (Fig. 4D). We measure the influence of each individual perturbation on each host gene through sensitivity analysis. We define the influence of perturbation *j* on gene *i* as 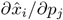 which can be calculated through the chain rule as

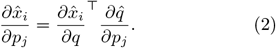

This measure of influence can be calculated for every sample, *x*, in the RNA-seq dataset and this is visualized in Fig. 4D (left column). Computing the influence of each perturbation, *p*_*j*_, on *araD*, we find that the arabinose concentration (Ara) has the most significant positive influence on expression. This is consistent with prior knowledge of Ara activation of the *araBAD* operon [51]. Similarly, we find that the model infers that *lacZ* is activated in the presence of IPTG, consistent with prior knowledge [51].

Using counterfactual (CF) experiments, our model infers that the change in wild-type expression between +Ara/-IPTG and -Ara/-IPTG can be explained by a simulated Ara dose and how much of the change in wild-type expression between -Ara/+IPTG and -Ara/-IPTG can be explained by a simulated IPTG dose. The CF experiment answers the question – what would the transcriptional profile have been if the strain was perturbed with a distinct perturbation vector *p*. We find that the model attributes the increase in expression in the two cases to Ara and IPTG induction (Fig 4D, right column), consistent with prior knowledge.

Using this measure of influence, we found that the expression of the synthetic cell state biomarkers (Fig. 3C) are impacted more than 87% of genes (3480 genes) in the *E. coli* genome with the biomarker impacted the most being impacted more than all but 3 genes – *lacZ, lacY, lacA* (Fig. 4E). The synthetic network influence score for gene *i* is calculated as

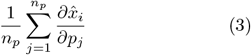

where *j* goes from 1 to *n*_*p*_ = 9 representing each perturbation *p*_*j*_ on the host from the synthetic gene expression of the 9 genes encoded on the NAND Circuit.

Our model quantitatively captures that kanamycin and chloramphenicol resistance markers significantly impact the synthetic cell state biomarkers (Fig. 4F). We also find that the basally active synthetic transcriptional regulators: TetR, LacI, LuxR, and AraC impact the host more than the transcriptional actuators PhlF and IcaR, which have little influence on synthetic cell state biomarkers. This may be due to the fact that the regulators are always expressed, while PhlF and IcaR are only expressed when IPTG or Ara are present, respectively.

In addition, we find that several synthetic cell state biomarkers (*fecCDE, yjiH, znuC, mepM*) are influenced only minorly by individual components of the synthetic gene network. One explanation for this is that these genes are influenced by combinations of perturbations rather than any single perturbation alone. In the next section, we use our model to test this hypothesis and show that phenotypes resulting from combinatorially influenced genes are generalizable.

### Synthetic cell state reveals antimicrobial resistance

We next sought to study the phenotypic impacts of the synthetic microbial cell state. Given that the expression of a significant number of genes in the *E. coli* genome were permanently dysregulated in the presence of a synthetic construct, it is expected that this results in unintended phenotypic consequences. We used our circuit-impact model to identify a subset of these genes for which no single perturbation results in significant dysregulation, the hypothesis being that these are genes which are impacted by synthetic constructs in general, rather than by what is being expressed on the construct. To test the generality of the phenotypes, we perform the following phenotypic screens in new strains in which the RNA-seq measurements were not performed.

We zoomed in on a highly-impacted gene, *mepM*, and used the circuit-impact model to explain the downregulation of the corresponding mRNA (Fig. 5A). We found that no single perturbation explains the dysregulation (Fig. 5B). We then performed the counterfactual experiment of setting combinations of the perturbations to zero to identify which set of genes are jointly responsible for the dysregulation. Setting the transcript levels of PhlF, IcaR, YFP, KanR, and CmR to zero results in a counterfactual *mepM* gene expression profile that largely recapitulates wild-type *mepM* transcript levels (Fig. 5C). This result indicates that the the downregulation of *mepM* may be a general synthetic microbe feature rather than one that is caused by an individual circuit component.

**Figure 5.**
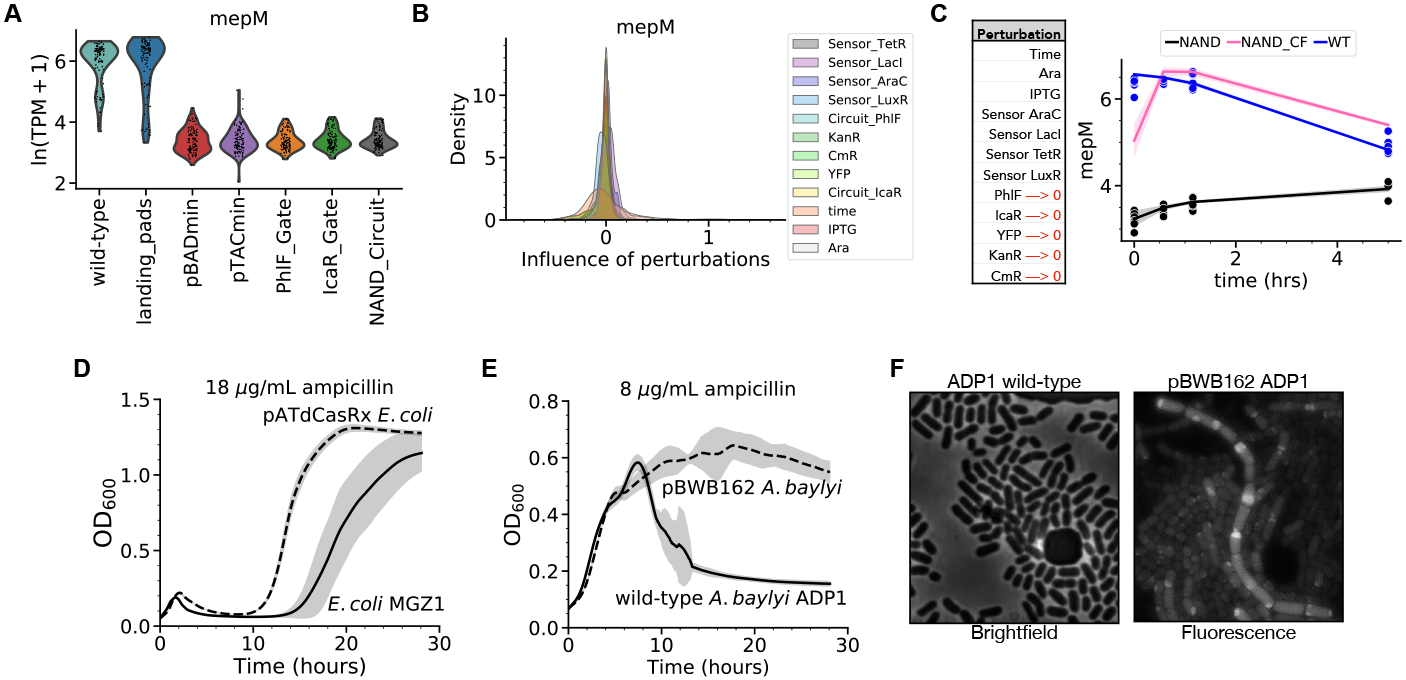
: Increased ampicillin resistance is a generalizable consequence of the synthetic cell state. **(A)** Distribution of the expression of *mepM* is shown for the wild-type strain and all genomic integration strains. Expression is quantified using ln(transcripts per million+1). **(B)** Kernel density estimates of the influence of synthetic gene perturbations on *mepM* as calculated through sensitivity analysis on the trained circuit-impact model. **(C)** The resulting *mepM* levels after performing the counterfactual experiment of setting a subset of synthetic gene levels to zero in the NAND Circuit strain (pink curve). Model predictions are visualized with solid lines while RNA-seq data are visualized with markers. The counterfactual result (NAND CF) is compared with the NAND Circuit expression and with wild-type (WT) expression. **(D)** Growth curves of *E. coli* MG1655 Z1 with and without the plasmid pATdCasRx subject to 18 *μ*g/mL ampicillin. **(E)** Growth curves of *A. baylyi* ADP1 with and without the plasmid pBWB162 subject to 8 *μ*g/mL ampicillin. **(F)** Brightfield and fluorescence microscopy images of wild-type ADP1 and pBWB162 ADP1 subject to 8 *μ*g/mL ampicillin, respectively.

Through high-throughput screens of cell envelope bioprocesses in *E. coli*, Babu et al. [52] showed that the lack of MepM results in impaired membrane invagination and cytokinesis. Based on their results, we hypothesized that the downregulation of *mepM* in synthetic strains should cause a decreased sensitivity to the *β*-lactam class of antibiotics.

The hypothesis we next test is: Do synthetic constructs induce a decrease in sensitivity to *β*-lactam antibiotics in engineered microbes? To test this, we challenged 5 engineered strains containing distinct genetic circuits with *β*-lactam antibiotics. First, we found that the introduction of the plasmid, pATdCasRx, containing a chloramphenicol resistance marker and a dCasRx-gRNA cassette in *E. coli* MG1655 Z1 (note that RNA-seq studies were conducted using MG1655) results in an increased resistance to ampicillin (Fig. 5D). We treated both the wild-type strain and the engineered strain with 18 *μ*g/mL of ampicillin which is above the known minimum inhibitory concentration for *E. coli* strains [53]. At this concentration of ampicillin, the engineered strain not only has a decreased lag time compared to the wild-type, but it also has a higher stationary phase cell density (Fig. 5B). We further tested the ampicillin resistance at a reduced temperature during growth and reduced concentration of ampicillin in the same engineered MGZ1 strain and another engineered strain which contains two plasmids. In both cases, the reduced ampicillin sensitivity was recapitu-lated (Supplementary Fig. 9A and B). Extending beyond ampicillin, we also find an increased resistance to the antibiotic cefazolin, another in the *β*-lactam class (Supplementary Fig. 9C). Testing for increased ampicillin resistance in the two plasmid-based NAND strains considered in this study, we find a decreased lag-time with respect to the wild-type but a lower stationary phase density.

To further test the generality of the reduced sensitivity to ampicillin across different strains and genetic components, we transformed another gram-negative strain, *Acinetobacter baylyi* ADP1, with the plasmid pBWB162 containing a kanamycin resistance marker, a *lacI* coding sequence, and an IPTG-inducible promoter driving the expression of *mCherry*. We found that the gram-negative ADP1 also shows an increased resistance to ampicillin in the presence of synthetic constructs. There is a drastic increase in the stationary phase cell density in engineered ADP1 strains compared to the wild-type (Fig 5E). In contrast to the previously considered strains, in the case of ADP1 in the absence of ampicillin, there is no reduction in growth rate relative to the wild-type strain. We find a minor increase in stationary phase cell density in the gram-positive strain *Bacillus subtilis* 168 subjected to ampicillin (Supplementary Fig. 10).

To connect the reduced ampicillin sensitivity with cell morphology and growth defects, we next imaged single cells under the microscope. We found that under treatment of ampicillin, ADP1 forms chains of daughter cells connected by uncleaved septa under induction of synthetic constructs (Fig. 5F). No chains of cells were visible in the wild-type strain. This is consistent with the results in [52], in which chains of daughter cells were apparent in *E. coli mepM* mutants (formerly known as *yebA*).

Purifed MepM is a peptidoglycan endopeptidase, essential and sufficient for the insertion of new glycan strands into the peptidoglycan mesh between the outer and inner cell membranes in growing gram-negative bacteria [54]. The splitting of daughter cells at the septa by peptidoglycan hydrolases such as MepM is required for cell division [55]. Babu et al. [52] discovered that *E. coli mepM* mutants show impaired membrane invagination and cytokinesis, resulting in what are chains of daughter cells connected by uncleaved septa. Seemingly, this same effect is reproduced here in *A. baylyi* ADP1 induced by synthetic constructs. Using BLAST homology searches, we were unable to identify *mepM* homologs with high confidence. The most similar coding sequence identified is the M23 family metallopeptidase (locus tag ACIAD RS15865) which contains a 312 bp region with a 46.2% similarity with *mepM*.

## Discussion

Through high-throughput RNA-seq analysis, we measured the transcriptional disruption imposed by synthetic genetic constructs integrated on the genome and plasmid-based constructs. Despite changing genetic contexts, as we introduce larger genetic circuits into the host, similar dysregulation patterns across strains were able to be identified and synthetic construct induced biomarkers were characterized. This suggested the existence of a synthetic microbial cell state that can be studied and exploited.

Through data-driven modeling of the transcriptional response to genetic perturbations, we quantified the impact of individual synthetic construct components on the host. Consistent with previous findings, the model identified that in a plasmid-based setting, the impact of antibiotic resistance genes on the host is significant. However, in the context of low-copy numbers with genomically-integrated constructs, the impact of transcriptional regulators outweighs the impact of the antibiotic resistance genes. Furthermore, the model infers that a majority of the synthetic cell state biomarkers are influenced by a combination of perturbations, rather than any single one. This suggested that the burden induced by the circuit on the biomarkers arise due to effects such as resource reallocation and can result in generalizable consequences.

The burden imposed by synthetic constructs, which is not always reflected in a reduction in growth rate, was shown to be generalizable to new strains and new constructs through the phenotype of increased resistance to antibiotics in the *β*-lactam class. This is a consequence of the downregulation of *mepM* (formerly known as *yebA*) in *E. coli* and hypothetically is due to a downregulation in peptidoglycan endopeptidases in *A. baylyi*.

Our modeling and data suggest that there are an enormous number of possible generalizable consequences of synthetic construct burden. Previous characterizations of synthetic construct burden were done in the context of fitness-burdened hosts, however our results in Figure 2A suggest that a burden on transcription is present even though growth profiles are comparable to the wild-type strain. The marked differences in housekeeping genes in Figures 2C and 4F, e.g., proteins associated with the outer membrane, the chaperone protein greB, biosynthesis genes from the wca and curli production cluster, suggest that circuit impact can occur from transcription and translational resource reallocation even if growth is not impacted. Further investigations should collect measurements that are not only probing transcription, but also epigenetic relationships and proteomic measurements could shed light on the mechanisms by which burden is imposed on the host.

## Methods

### Strains and plasmid engineering

The strain used in this study of the transcriptional response to synthetic constructs is *E. coli* MG1655 (NCBI genome sequence U00096.3; [56]). Cell growth for cloning and plasmid extraction were performed as in [57] (see Materials and Methods section “Strains and media”) unless otherwise noted.

Two other strains used in this study, namely *E. coli* MGZ1 [58] and *A. baylyi ADP1* (NCBI genome sequence NC 005966.1; [59]) have distinct growth and cloning protocols. Both were transformed with plasmids containing the medium-copy pBAV1K backbone [60].

Chemically competent MGZ1 were prepared using TSS Buffer and the protocol from [61] was followed. Transformation of MGZ1 was performed as follows. Chemically competent MGZ1 cells were thawed on ice and 1 *μ*L of prepped plasmid was was mixed gently with a pipette. The cells were incubated on ice for 15 minutes followed by incubation at 42^°^C for 30 seconds. The cells were incubated on ice for 2 minutes. 1 mL of LB media was added at room temperature followed by incubation and shaking at 37^°^C for 1 hour. Transformed cells were plated on appropriate antibiotic LB agar plates and grown overnight at 37^°^C.

Naturally competent ADP1 was grown overnight in LB media in a shaking incubator at 30^°^C. 70 *μ*L of cells along with 100 ng of plasmid DNA were added to 1 mL of fresh LB. The cells were incubated and shaken for 3 hours followed by plating on appropriate antibiotic selection LB agar plates.

### Construction of genomic landing pads and insertion of payloads

Genomically-integrated landing pad strains of *E. coli* MG1655 constructed and verified in [57] were used in this study for payload integration of the genetic NAND gates (For details, we refer to Materials and Methods section “Construction of genomic landing pads” and section “Insertion of payloads into landing pads”).

Briefly, in [57] three landing pads were inserted into the genome of *E. coli* MG1655. Two landing pads were introduced sequentially using *λ*-RED recombineering [62]. A third landing pad was constructed and inserted using a site-specific mini-Tn7 transposase [63]. After the genomic insertion of landing pads, each landing pad contained a unique antibiotic resistance marker (chloramphenicol, kanamycin, and tettracycline for Landing Pads #1, #2, and #3, respectively) located between a pair of unidirectional flippase recognition target (FRT) sites.

For insertion of payloads into the landing pads, first cotransformation of the empty landing pads strain with a plas-mid encoding three integrases [57] and a plasmid containing the DNA payloads. Electrocompetent cell preparation and transformation was performed as in [57] and integration was confirmed with colony PCR by amplifying regions containing integrated constructs and adjacent genomic DNA.

### RNA sequencing and data processing

Wild-type and engineered *E. coli* MG1655 were cultured overnight in M9 Media consisting of 1X M9 media salts, 0.1 mM CaCl_2_, 1X trace salts, 1 mM MgSO_4_, 0.05 mM FeCl_3_, 0.1 mM C_6_H_8_O_7_, 0.2% Casamino Acids, and 0.4% Glucose.

Glycerol stocks were inoculated into M9 media in shake flasks, and the culture was grown overnight for 18h at 30^°^C and 1000 rpm. The following day, cultures were diluted to OD 0.1 in fresh M9 media and grown in 96-well plates under the same conditions for 3 hours. For induction, cells were diluted a second time to OD 0.05 in the presence of inducers. Plates were incubated at 37^°^C and 1000 rpm for 5 hours, 6.5 hours, 8 hours, and 18 hours and cultured cells were harvested and fixed with RNAprotect (Qiagen 76506).

Total RNA was extracted using Magjet RNA extraction kit (Thermo Fisher) according to the manufacturer’s instructions. RNA quality was assessed using a Tapestation (Agilent). KAPA RNA Hyperprep Kit (Roche 08098140702) was used for ribosomal RNA depletion and Illumina compatible library preparation. The prepared library was loaded on an Illumina sequencer to generate 150 base pair paired end reads.

The raw RNA sequencing data were trimmed and quality filtered with Trimmomatic (v0.36) [64]. The trimmed reads were aligned to the *E. coli* genome with bwa (v0.7.17) [65]. Gene-level quantification of counts was performed using the featureCounts function of Rsubread (v1.34.4) [66].

We followed [26] and used three metrics to measure a sample’s quality: 1) number of reads mapped must be greater than or equal to 5 × 10^5^, 2) count of all annotated genes must be greater than 5 × 10^5^, and 3) replicates must have correlation greater than 0.9. If a sample did not pass any of these three metrics, than the sample was discarded and not used for down-stream analysis.

### Differential expression, gene ontology analysis

The Python package for ‘omics analysis scanpy [67] was used to perform differential expression analysis. Specifically, log-transformed transcripts per million were used with a Wilcoxon rank-sum test were used to test for differential expression between wild-type and NAND Circuit strains. The p-values associated with the log fold changes were corrected for multiple testing using the Benjamini-Hochberg procedure.

Gene ontology enrichment analyses [37] were performed using PANTHER [68]. All identified gene sets were tested for statistical significance using Fisher’s exact test and gene sets were only considered significant if the false discovery rate did not exceed 5%.

The database RegulonDB [69] was used to identify transcription factors and sigma-factors for differentially expressed genes and syntehtic cell state biomarkers.

### Predicting optical density from RNA-seq measurements

RNA-sequencing measurements at all sampled time points (5, 6.5, 8, and 18 hours) and biological replicates (eight) were used to build a linear model to predict the optical density (OD_600_) at 18 hours after growth. In order to prevent overfitting to the data and assess prediction variance across subsamples of the data, we divided the data over 192 folds stratified by biological replicates. The OD observations in a fold were averaged across biological replicates, *y*^*s*^, from each strain, *s* for *s* ∈ {wild-type, pBADmin, pTACmin, PhlF Gate, IcaR Gate, NAND}, and concatenated into the vector

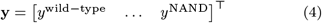

with **y** ∈ ℝ^6^. We then selected *n* genes from the transcriptome and formed the *n* × 6 data matrix **X** with *x*_*is*_ computed as the average expression of gene *x*_*i*_ in strain *s* in a fold. By taking the average of the expression over time, we are implicitly give equal weight to the expression at each time point. We then constructed a linear model of the form

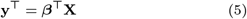

where *β* ∈ ℝ^n^ is a vector of coefficients that weigh the contribution of each gene towards prediction of the OD observations. We then minimize the squared error between the left-hand side and right-hand side, 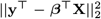 and the solution is obtained by taking the gradient with respect to *β* to obtain the least squares solution:

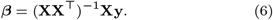

We calculate the goodness of fit of the parameters in *β* through the coefficient of determination, *R*^2^, and through the Pearson correlation between the mean predicted OD and the mean measured OD. The mean of the predictions are taken over the 192 folds while the mean of the measurements are taken over biological replicates. These averages and standard deviations were reported in the Results section and used within figures.

### Classifying synthetic-construct-induced biomarker genes

The transcriptional profiles were assigned labels: 0 for a wild-type strain and 1 for an engineered strain. Data were z-scored and a binary logistic regression model was trained on a training set of data consisting of 50% of all samples. The training set consisted of 503 samples of which 49 samples were from the wild-type class and the remaining belonging to the synthetic class. The test set, containing 63 wild-type samples and 441 synthetic samples, are used to measure the empirical performance of the classifier.

L2 regularization was used to prevent the model from over-fitting to the training data. A 5-fold cross-validation was performed on the training set to determine the optimal L2 regularization weight. Once the L2 regularization weight was identified through cross-validation, the logistic regression model was then re-trained on all of the training data. Testing the model on the test set, we found that it accurately classified all 504 test points.

The magnitude of the model coefficients were then used to assign feature importances to each of the genes; the higher the magnitude of the coefficient, the more important a gene is for classificatiion. This was validated through training a sequence of logistic regression classifiers on only important features and through comparison to classifiers trained with random features.

### Circuit-impact model

Our dataset consists of the triplet of measurements 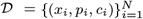, where each *x*_*i*_ ∈ ℝ^g^ denotes the gene expression of all *g* genes from sample *i, p*_*i*_ ∈ ℝ^m^ describes *m* individual, continuous perturbations applied to the cells in sample *i*, and *c* ∈ ℝ^n^ describes any discrete covariate that is to be modeled. For example, discrete covariates can be a batch, patient, species, and for the circuit-impact model it is describing whether and how the synthetic construct is integrated into the host. More specifically, the perturbations in the circuit-impact model are,

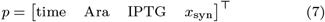

where *x*_syn_ is the vector of synthetic gene expressions. If *p*_*ij*_ = 0, then perturbation *j* was not applied to sample *i*. Similarly, the covariates in the circuit-impact model are,

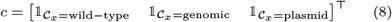

where 𝟙 denotes and C is the set representing whether a sample *x*_*i*_ is from the wild-type strain, a genomically-integrated strain, or a plasmid-based strain.

Given the dataset, our task is to model the impact that perturbations, *p*, and covariates, *c*, have on the host gene expression *x*. We start by assuming that there exists a latent (hidden) state, *z*_*i*_ ∈ ℝ^l^, for each gene expression vector *x*_*i*_, such that *z*_*i*_ contains no information about the corresponding perturbation, *p*_*i*_, or covariates, *c*_*i*_. We call this the perturbation-free latent state. Simultaneously, we assume that there are latent representations of the perturbations, *q*, and covariates, *d*, such that they impact the perturbation-free latent state, *z*, additively, producing a perturbed latent state, 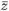. This assumes that the perturbation-free latent state is independent of the latent perturbations and latent covariates. The perturbed latent state is formed directly as

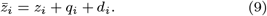

### Deep learning architecture and training of the circuit-impact model

The architecture of the circuit-impact model consists of three encoders, one for each of the triplets in D, i.e. a gene expression encoder, *F* : ℝ^g^ → ℝ^ml^, a perturbation encoder, *G* : ℝ^m^ → ℝ^l^, and a covariate encoder, *H* : ℝ^n^ → ℝ^l^, such that

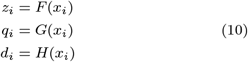

which can then be used to form the perturbed latent state 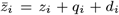. The perturbed latent state is then transformed back to gene-space through the decoder 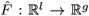 to produce, 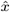, a reconstruction of the original gene expression vector. Similarly, *q* and *d* are decoded into reconstructions 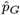 and 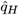, respectively.

To facilitate the encoder *F* to map gene expression vectors to a perturbation-free latent state, we employ two techniques: 1) enforce that *z*_*i*_ ∼ N(0, *I*), and 2) train discriminators, *D*_*p*_ and *D*_*c*_ to predict the perturbations, *p*_*i*_, and covariates, *c*_*i*_, given *z*_*i*_, respectively. The normally distributed latent states encourage a specific geometric structure, but inherently cannot prevent clusters that group strains together from forming. This is where the discriminator architecture facilitates the latent state to be perturbation free by penalizing the encoder losses for good discrimination. This form of adversarial training to disentangle factors of variation from input features follows from [28].

The mean squared error is used to calculate the reconstruction loss between *x, p* and their respective decoder outputs 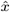, 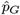:

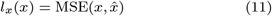

where *B* is the batch size and *y* and 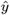 are the considered inputs to be reconstructed and the network reconstruction, respectively. The covariate autoencoder loss and the covariate discriminator, *D*_*c*_, loss is calculated as the cross-entropy between the output of the covariate decoder (*H*^−1^) and the output of *D*_*c*_ and the true covariates, respectively:

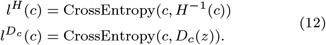

The mean squared error is also used to calculate the discriminator loss for the continuous perturbation discriminator (*D*_*p*_):

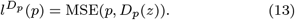

Finally, the Kullback-Leibler (*KL*) divergence between the perturbation-free latent state distribution and a standard normal prior is calculated as the latent space distribution penalty:

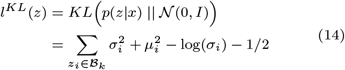

where ℬ_∥_ is the set of samples for a single batch of data *k, σ* is the vector of variances of the perturbation-free latent state *z*, and *μ* is the vector of means of the perturbation-free latent state.

The circuit-impact model is trained in steps wherein the parameters of the discriminators, encoders, and decoders are sequentially optimized. For a batch ℬ_*k*_ consisting of 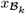, 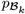, and 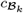, training takes places in three steps:

1. Minimize 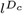 and 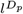by updating the parameters of D_c_and D_p_, respectively;
2. Minimize 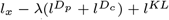 by updating the parameters of *F*, 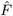, *G*, and *H*.
3. Minimize *l*_*p*_ and *l*^*H*^ by updating the parameters of *G* and *H*, respectively.

We use the Adadelta optimizer [70] for parameter optimization and implement the model in Pytorch.

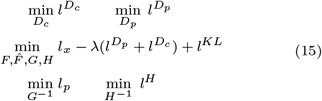

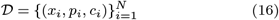

### Hyperparameter Selection and Training

The hyperparameters we considered during training of the circuit-impact model were the penalty coefficient, *λ*, for good discriminator performance, the encoder and decoder learning rates, L2 regularization weight on network parameters, the batch size, the dimension of the latent state *l*, and the dimension of hidden layers in the network. 50 models were trained for randomly chosen hyperparameters. The following table depicts the distribution over which each hyperparameter was selected from are given in Table 1.

**Table 1:** Random search over hyperparameters.

| Hyperparameter | Random Search |
| --- | --- |
| $\lambda$ | Uniform($10^{-4}, 2^{-3}$) |
| learning rate | Uniform(1, 2.1) |
| L2 regularization | Uniform($0, 10^{-6}$) |
| batch size | $2^b$, $b = 3, \dots, 9$ |
| latent state dim. $l$ | $\{5, 6, 7, 8, 9, 10, 11, 12\}$ |
| hidden layer dimensions | $\{128, 256, 512\}$ |

The dataset was stratified by strain and cross-validation was performed by training the 50 models with randomly selected hyperparameters on each of the stratified validation datasets. We used *R*^2^ as the metric of accuracy to choose the hyperparameters which resulted in a model with highest average accuracy across the validation datasets. We then trained the final model on all strains using these hyperparameters.

### Calculating influence of perturbations on host genes

Once the model is trained, we can calculate the derivate of the output of the network gene encoder, 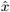, with respect to the perturbation *p* to obtain a measure of sensitivity of each host gene to each perturbation. Specifically, for a sample *x*_*i*_ corresponding to perturbations *p*_*i*_ and covariates *c*_*i*_, to calculate the influence of perturbation j, *p*_*ij*_, on gene *k*, 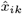, we compute 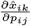. We can perform this operation simultaneously for all N datapoints in 𝒟, producing a tensor, *J* ∈ ℝ;^*N* ×*g*×*m*^, where *J*_*ijk*_describes the influence of perturbation *k* on gene *j* in sample *i*. **Cell imaging with CellASIC** *Acinetobacter baylyi* ADP1 from a glycerol stock was recovered overnight in LB and passaged to log-phase to reach a target OD of 0.3. Cells were subsequently diluted, per manufacturer recommendations, to OD 0.05, or 5 × 10^7^ cells per milliliter and then loaded into an inlet valve in the CellASIC plate. The cells were then primed and injected into the chamber using 0.25 kPa of pressure, suspended in standard LB media (Teknova L8000). The Cell ASIC plate was mounted on an automated Advanced Scientific Imaging microscopy stage for planar control within an Olympus IX83 epifluorescence microscope system. The entire plate was incubated at 30°C in a Tokai incubation chamber mounted on the stage of the Olympus IX83 frame. Cells were imaged with brightfield illumination and fluorescence illumination (580 nm excitation and 610 nm emission) every 12.5 minutes for 15 hours. A media of LB and ampicillin at 8 *μ*g/mL was injected after 3 hours of cell growth in the chamber.

## Data availability

Processed RNA-seq data used in this study are available at: https://github.com/AqibHasnain/circuit-state. The RNA sequencing data generated in this study have been deposited in the GEO database under accession code GSE206047

## Code availability

Code to reproduce all figures in this study is available at: https://github.com/AqibHasnain/circuit-state.

## Author contributions

AH, AEB, YD, CAV, and EY conceived of the study. AEB, YD, CAV, and EY designed experiments. AEB and YP constructed the synthetic gene circuits. DB and PM performed RNA-seq experiments. JU processed RNA-seq measurements. AH performed data analysis and predictive modeling with feed-back from SB and EY. AH and LR performed growth and microscopy experiments. AH wrote the manuscript with inputs from all authors.

## Acknowledgments

This work was supported by the US Defense Advanced Research Projects Agency (DARPA) Synergistic Discovery and Design program (SD2). We thank Alec Taylor from University of California Santa Barbara, Department of Mechanical Engineering for providing the dCasRX plasmid.

## Notes

https://www.ncbi.nlm.nih.gov/geo/query/acc.cgi?acc=GSE206047

### Competing Interest Statement

The authors have declared no competing interest.

