## Supplementary Material for "Disentangling gene expression burden identifies generalizable phenotypes induced by synthetic gene networks"

Aqib Hasnain et al.

#### 1 Supplementary Figures

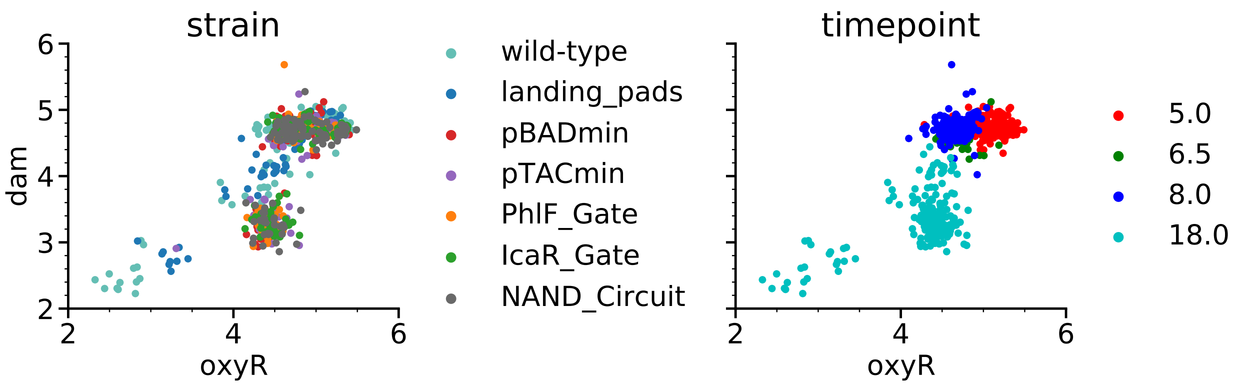


Figure 1: Expression of the gene *oxyR* encoding the repressor OxyR and the DNA adenine methyltransferase coding gene *dam* colored by strain (left) and time point (right).

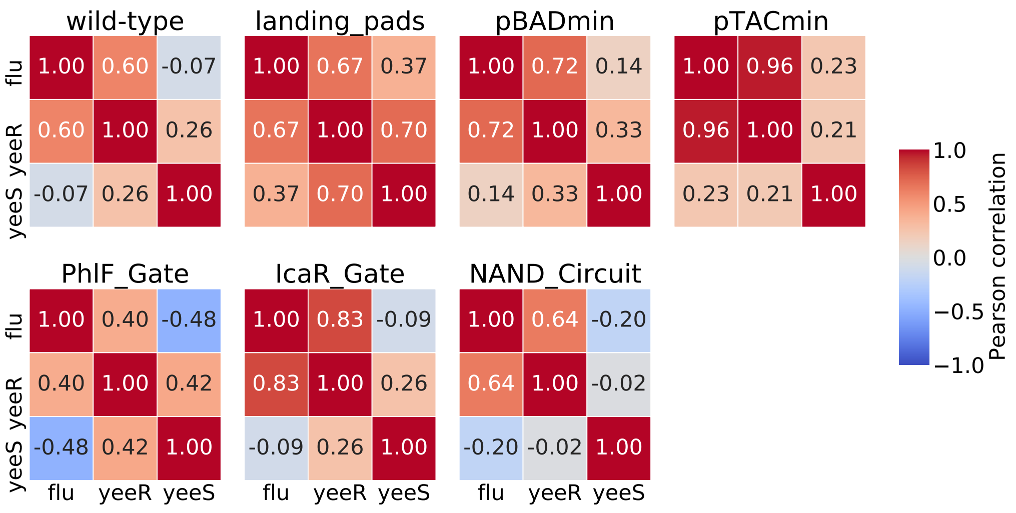


Figure 2: Pearson correlations between the RNA-seq measurements of the three genes, *flu*, *yeeR*, and *yeeS*, used to characterize the strain-to-strain fitness variability.

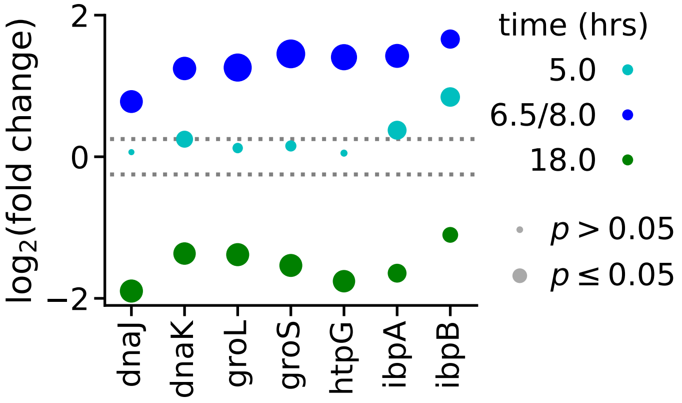


Figure 3: Fold change of the synthetic-construct-induced biomarkers from Ceroni *et al.* comparing the NAND circuit to the wild-type condition. The markers size corresponds to the multiple-testing corrected p-value: smaller markers have higher p-values.

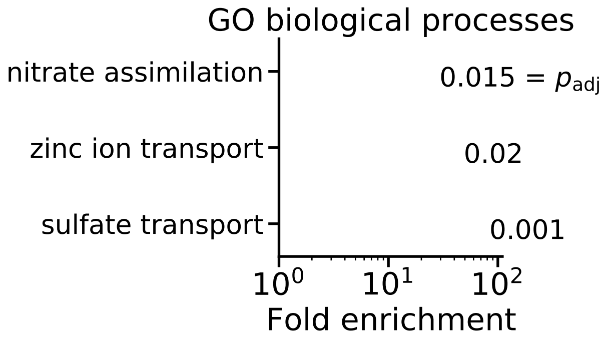


Figure 4: Overenriched gene ontologies for the 33 genes classified to be synthetic cell state biomarkers.

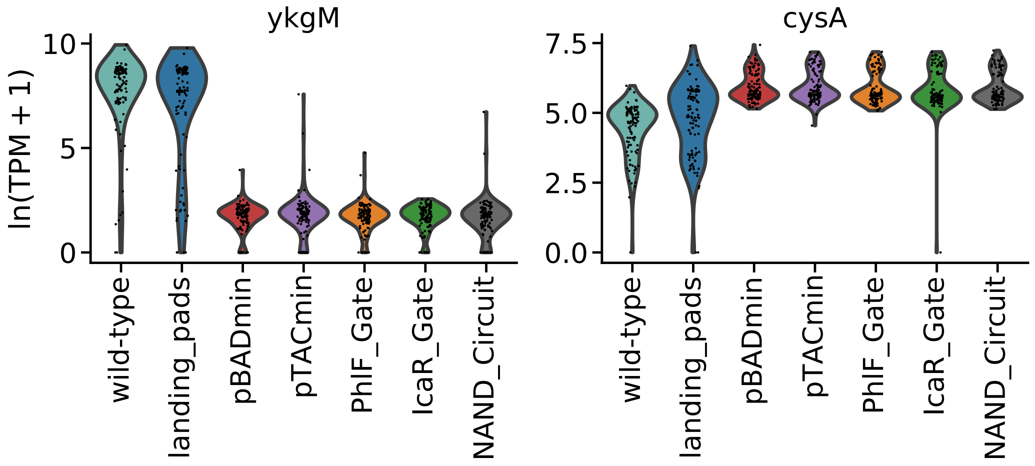


Figure 5: Intra-strain distributions of the synthetic cell state genes with lowest fold change and highest fold change with respect to the wild-type strain.

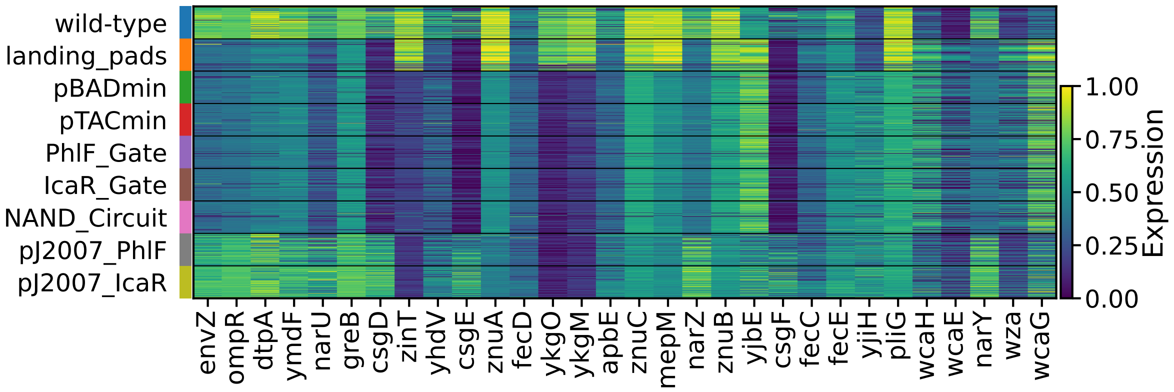


Figure 6: Expression (minmax normalized across all conditions) for synthetic cell state genes grouped by strain and including the two plasmid strains.

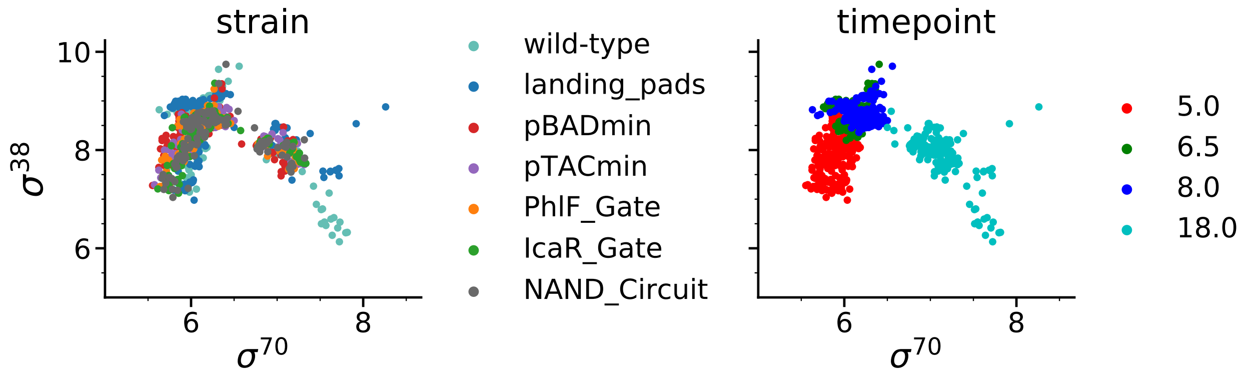


Figure 7: Expression of the gene *rpoS* encoding the sigma factor $\sigma^{70}$ and the gene *rpoD* encoding the sigma factor $\sigma^{38}$ colored by strain (left) and time point (right).

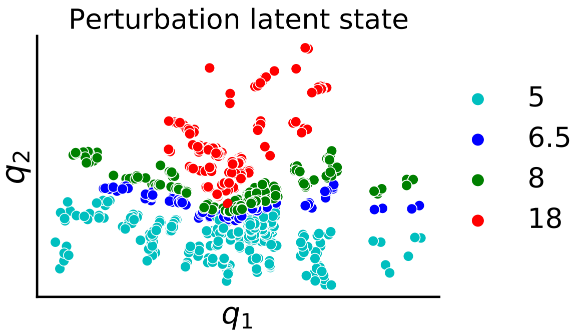


Figure 8: The circuit-impact model's perturbation latent state. The markers are colored by time point, showing that the second dimension of the latent state is largely capturing time, a significant perturbation to the host transcriptome.

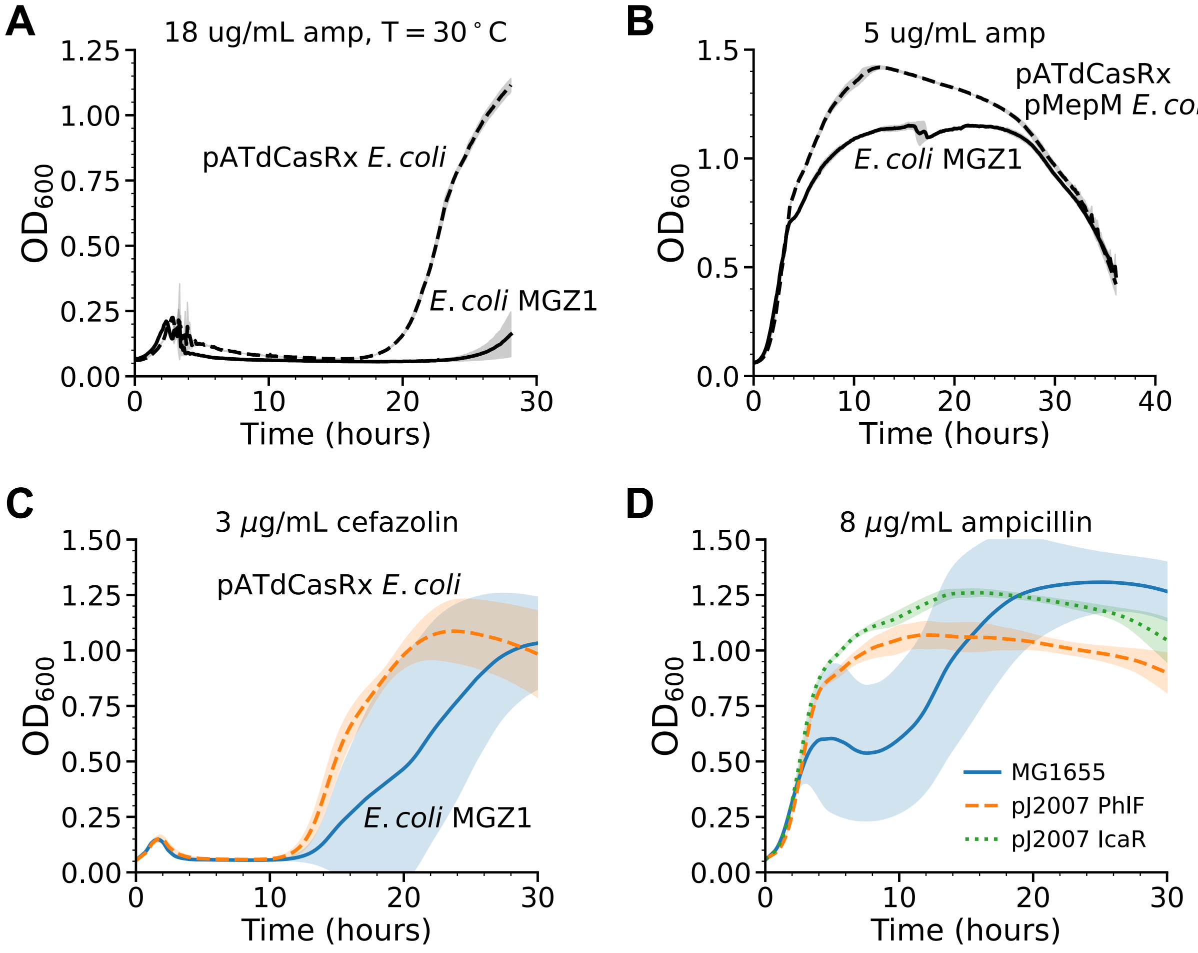


Figure 9: Growth curves comparisons for E. coli strains treated with $\beta$-lactam antibiotics. **(A)** Growth curves taken at 30°C and 18 $\mu$g/mL ampicillin. **(B)** Growth curves taken at the standard growth temperature of 37°C and 5 $\mu$g/mL ampicillin. **(C)** Growth curves taken at 37°C and 3 $\mu$g/mL cefazolin, another antibiotic in the $\beta$-lactam claass of antibiotics. **(D)** Growth curves for the two plasmid strains considered in this study and the corresponding wild-type strain, MG1655, subject to 8 $\mu$g/mL.

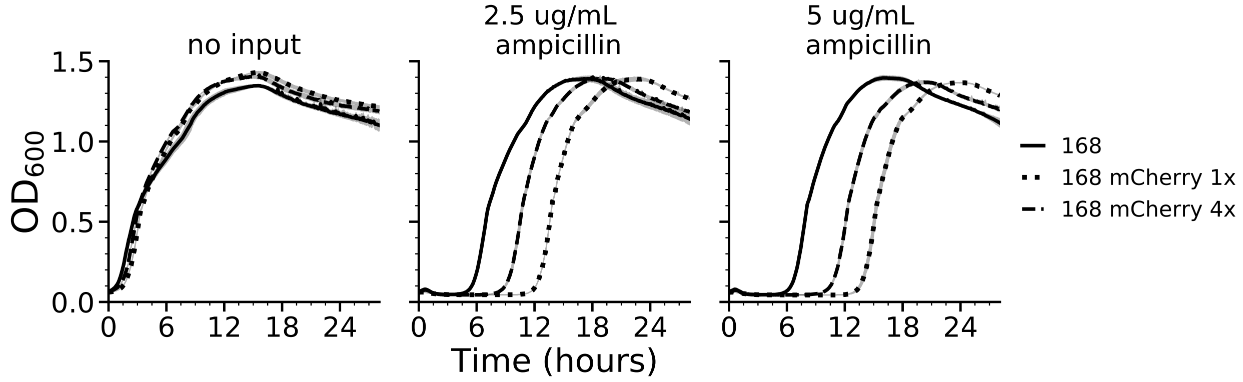


Figure 10: *Bacillus subtilis* 168 growth curves for the wild-type strain, the strain with 1 copy of mCherry on the genome, and the strain with 4 copies of mCherry on the genome. The ampicillin concentrations tested are 0, 2.5, and 5 $\mu$g/mL and resulting growth curves are plotted in the left, middle, and right panel, respectively.

### 2 Supplementary Tables

| regulator | regulated | effect |
|---|---|---|
| Sigma24 | wza | + |
| Sigma38 | csgD | + |
| Sigma38 | csgE | + |
| Sigma38 | csgF | + |
| Sigma38 | narU | + |
| Sigma38 | znuA | + |
| Sigma70 | zinT | + |
| Sigma70 | ykgO | + |
| Sigma70 | ykgM | + |
| Sigma70 | yjbE | + |
| Sigma70 | wza | + |
| Sigma70 | ompR | + |
| Sigma70 | dtpA | + |
| Sigma70 | envZ | + |
| Sigma70 | znuB | + |
| Sigma70 | csgF | + |
| Sigma70 | csgE | + |
| Sigma70 | csgD | + |
| Sigma70 | narU | + |
| Sigma70 | znuC | + |

Table 1: Sigma factor-gene regulation of the synthetic cell state biomarkers.

| regulator | regulated | effect |
|---|---|---|
| BasR | csgD | + |
| BasR | csgE | + |
| BasR | csgF | + |
| CpxR | csgF | - |
| CpxR | csgE | - |
| CpxR | csgD | - |
| Cra | csgD | + |
| Cra | csgE | + |
| Cra | csgF | + |
| CsgD | csgD | + |
| CsgD | csgF | + |
| CsgD | csgE | + |
| FliZ | csgD | - |
| FliZ | csgE | - |
| FliZ | csgF | - |
| Fur | zinT | + |
| Fur | fecD | - |
| Fur | fecC | - |
| Fur | fecE | - |
| GadX | dtpA | - |
| HypT | fecE | - |
| HypT | fecD | - |
| HypT | fecC | - |
| MlrA | csgD | + |
| MlrA | csgE | + |
| MlrA | csgF | + |
| MqsA | csgD | - |
| MqsA | csgF | - |
| MqsA | csgE | - |
| Nac | fecE | NaN |
| Nac | fecC | NaN |
| Nac | fecD | NaN |

| regulator | regulated | effect |
|---|---|---|
| OmpR | csgF | + |
| OmpR | dtpA | + |
| OmpR | csgD | + |
| OmpR | csgE | + |
| OxyR | znuA | + |
| OxyR | znuB | + |
| OxyR | zinT | + |
| OxyR | znuC | + |
| PdhR | fecC | + |
| PdhR | fecE | + |
| PdhR | fecD | + |
| RcdA | csgD | + |
| RcdA | csgE | + |
| RcdA | csgF | + |
| RcsB | csgD | - |
| RcsB | csgE | - |
| RcsB | csgF | - |
| RcsB | yjbE | + |
| RcsB | wza | + |
| RstA | csgD | - |
| RstA | csgE | - |
| RstA | csgF | - |
| SoxS | zinT | + |
| Zur | znuB | - |
| Zur | znuA | - |
| Zur | znuC | - |
| Zur | zinT | - |
| Zur | ykgO | - |
| Zur | ykgM | - |

Table 2: TF-gene regulation of the synthetic cell state biomarkers.
